# Realistic simulation of time-course measurements in systems biology

**DOI:** 10.1101/2023.01.05.522854

**Authors:** Janine Egert, Clemens Kreutz

## Abstract

**Motivation:** In systems biology, the analysis of complex nonlinear systems faces many methodological challenges. However, the performance evaluation of competing methods is limited by the small amount of publicly available data from biological experiments. Therefore, simulation studies with a realistic representation of the data are a promising alternative and bring the advantage of knowing the ground truth.

**Results:** We present an approach for designing a realistic simulation study. Based on 19 published systems biology models with experimental data, we assess typical measurement characteristics such as observables, observation type, measurement error, and observation times. For the latter, we estimate typical time features by fitting a transient response function. We demonstrate the approach on a meal model of the glucose insulin system, a mitogen-activated protein-kinase cascade and a model for the epidermal growth factor signaling. The performance of the realistic design is validated on 9 systems biology models in terms of optimization, integration and identifiability. For any dynamic model downloaded from an online database, our algorithm analyzes the model dynamics and specifies a realistic experimental design. The approach is specifically suited for systematic benchmarking of methods for timecourse data in the context of systems biology. In particular, various application settings such as number of parameters, initial conditions, error model etc. can be tested.

**Availability:** The approach is implemented in the MATLAB-based modelling toolbox Data2Dynamics and available at https://github.com/Data2Dynamics/d2d.

## 1 Introduction

Systems biology deals with the mathematical analysis of complex biological processes and networks, often by means of large nonlinear systems. Therefore, many computational issues arise and large differences in performance between different application settings have been found [9, 19, 26, 29, 33, 34, 36]. Therefore, an important task is to evaluate the performance of methods and algorithms in both experimental applications and simulation studies.

Databases such as the BioModels database [22] offer a wide range of published biological models and provide equations for the dynamic variables of a desired biological system typically in the SBML format [16]. However, the BioModels database does not provide experimental data and how the dynamic equations are linked to the data. To fill this gap, there are attempts to assemble systems biology models together with the underlying experimental data in a benchmark repository [13, 37].

Although simulation studies can not replace experimental data, they are a valuable tool for evaluating different methods and algorithms used for systems analysis. In contrast to experimental data, they are advantageous in that an unlimited number of measurement and analysis repetitions can be simulated. Since the underlying truth of the system is known, additional statistical quantities, such as sensitivity and specificity, can be assessed from simulated data.

Here, we present an approach to create realistic simulations of any dynamic system and to provide an experimental design, including a reasonable reflection of observables, observation time points, and a realistic data set with measurement errors. Our results are based on the experimental design of 19 biological models of a benchmark repository [13] and can be applied to a variety of dynamic models e.g. from the BioModels database [22]. While typical proper-ties of the experimental design are provided by the 19 published published models, the dynamics are defined by the desired model.

## 2 Materials and method

This work focuses on biological processes that can be modeled by ordinary differential equations (ODE)

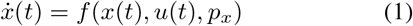

where *x* denotes the states of the biological system, *p* its parameters, and *u* the input function representing a stimulus of the system. Function *f* translates the interaction map of the biological processes into an ODE system. In a biological context, the states *x* correspond to molecular concentrations and represent the compartments of interest of the biological system. Because not all states *x* of a system can be measured in an experiment due to biological limitations or cost, often only data from a subset of states are collected. These measured states are defined by the experimenter prior to conducting the experiment and are referred to as observables

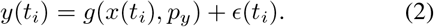

The function *g* links the dynamic states *x* with the measured values of the observable *y* and its parameters *p_y_*, and may include a logarithmic transformation. The additive term *ϵ* represents the measurement noise.

### 2.1 Observables

Typical observation functions *g* are

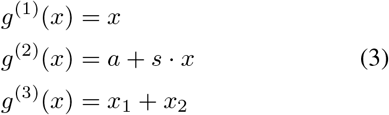

Here, *g*^(1)^ represents a direct measurement of the observable, *g*^(2)^ a relative measurement with an offset *a* and a scaling factor *s,* and *g*^(3)^ a combined measurement of two states *x*_1_ and *x*_2_, which in the following is also called a compound measurement. Common measurement techniques in biological applications for measurement type *g*^(2)^ are e.g. Western-blot, PCR, sequencing and proteomics experiments, and for *g*^(3)^ e.g. ligand-protein binding without discrimination of phosphorylation status. The measurement technique is chosen by the experimenter depending on measurability, accuracy, and cost of the experiment.

### 2.2 System dynamics

An essential part of setting up a realistic design is the choice of observation time points. Too short time duration does not capture the long-term dynamics, and larger time duration or larger intervals between the measurement points do not properly capture the fast dynamics. To define a reasonable set of observation time points which fit the model dynamics properly, the time scale of the desired model has to be estimated. Here, the time scale of the model dynamics is given by the transient function

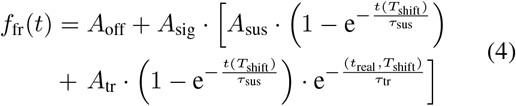

where *A*_off_ is an additional offset, *A*_sig_ is a conversion prefactor which can assume 1 or −1, *A*_sus_ and *A*_tr_ are the amplitudes of the sustained increase and the transient peak, and *τ*_sus_ and *τ*_tr_ are the time parameters of the transient function indicating the velocity of the dynamics [21]. Because there are cases where the dynamics start with a delay, the time predictor is shifted by a nonlinear transformation [21]

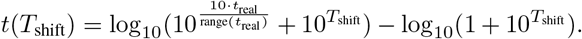

### 2.3 Realistic simulation

In this paper, we present an approach for a simulation of the experimental design and data for systems biology models. Biochemical reaction networks which can be modeled using transient dynamics are selected from the Data2Dynamics [28] examples folder, e.g. no oscillations are included. To this end, 19 biological pathway models [1, 2, 3, 4, 5, 6, 7, 10, 11, 14, 17, 24, 25, 27, 12, 31, 32, 35, 38] are analyzed for their experimental design. The analysis of the 19 benchmark experiments as well as the implementation of the realistic simulation are described in the following section. The main steps of the implemented algorithm are shown in Figure 1 and explained in more detail below.

**Figure 1:**
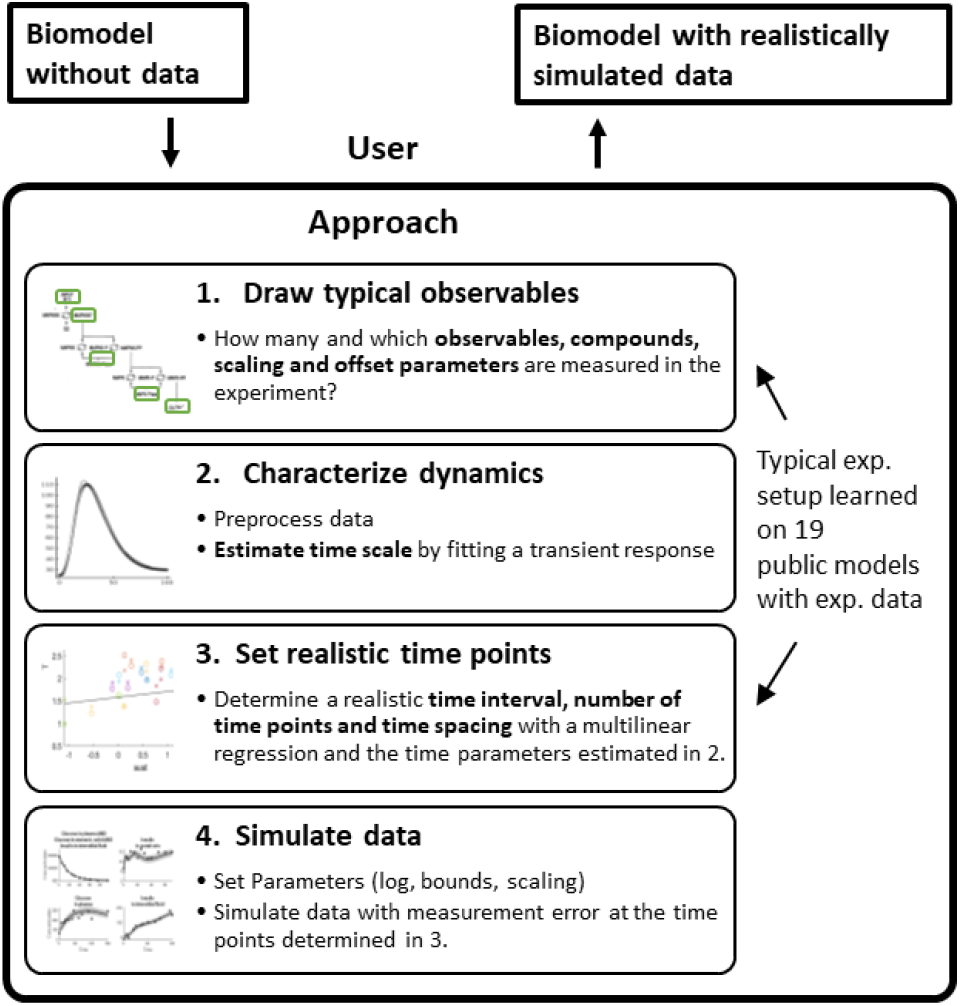
Main steps for setting up a realistic simulation for a desired model.

#### 2.3.1 Realistic observables and observation functions

The number of observables, relative observation functions and compound measurements of the 19 published models are depicted in Table 1. In these 19 experiments, on average 50% of the dynamic states are measured, of which 40% are measured directly (*g*^(1)^), 40% with a scaling factor (*g*^(2)^) and 20% contribute to a compound measurement (*g*^(3)^). In order to guarantee a realistic simulation setup, the number of observables and their observation function are drawn by a binomial draw from Table 1 Columns 4-8, and are then randomly assigned to the dynamic state variables.

**Table 1:** Frequency of observables and their observation type for the 19 published models.

| Model | # Species | # Obs | $\frac{\# \text{ Obs}}{\# \text{ Species}}$<br>[%] | $\frac{\# \text{ Compound}}{\# \text{ Obs}}$<br>[%] | # Species<br>in Compound | $\frac{\# \text{ Scaling}}{\# \text{ Obs}}$<br>[%] | $\frac{\# \text{ Offset}}{\# \text{ Scaling}}$<br>[%] | Ref. |
| --- | --- | --- | --- | --- | --- | --- | --- | --- |
| Alkan | 36 | 12 | 33 | 0 | 0 | 83 | 0 | [1] |
| Bachmann | 25 | 20 | 80 | 5 | 2 | 50 | 50 | [2] |
| Becker | 6 | 4 | 67 | 0 | 0 | 25 | 100 | [3] |
| Boehm | 8 | 3 | 38 | 100 | 2 | 0 | 0 | [4] |
| Brannmark | 9 | 3 | 33 | 33 | 2 | 0 | 0 | [5] |
| Bruno | 7 | 6 | 86 | 0 | 0 | 0 | 0 | [6] |
| Crauste | 5 | 4 | 80 | 0 | 0 | 0 | 0 | [7] |
| Fiedler | 6 | 2 | 33 | 0 | 0 | 0 | 0 | [10] |
| Fujita | 9 | 3 | 33 | 0 | 0 | 100 | 0 | [11] |
| Hass | 61 | 19 | 31 | 47 | 10.3 | 79 | 93 | [14] |
| Isensee | 25 | 3 | 12 | 0 | 0 | 67 | 0 | [17] |
| Lucarelli | 65 | 33 | 51 | 61 | 5.3 | 0 | 0 | [24] |
| Merkle | 23 | 22 | 96 | 5 | 2 | 50 | 82 | [25] |
| Raia | 14 | 8 | 57 | 13 | 3 | 63 | 0 | [27] |
| Reelin | 9 | 6 | 67 | 100 | 5 | 100 | 100 | [12] |
| Schwen | 11 | 4 | 36 | 100 | 2.8 | 50 | 100 | [31] |
| Sobotta | 35 | 18 | 51 | 0 | 0 | 67 | 100 | [32] |
| Swameye | 14 | 3 | 21 | 0 | 0 | 67 | 0 | [35] |
| Zheng | 15 | 15 | 100 | 0 | 0 | 0 | 0 | [38] |
| $\mu \pm \text{sd}$ | $20 \pm 20$ | $10 \pm 9$ | $50 \pm 30$ | $20 \pm 40$ | $2 \pm 3$ | $40 \pm 40$ | $20 \pm 30$ | |

#### 2.3.2 Observation time points

The choice of observation time points, i.e. the time duration and time spacing, play an important role in creating a realistic design. We observed that experimentally chosen time spacing in the published models commonly fall between linearly and exponentially distributed time intervals. Thus, we estimate the spacing of time points as:

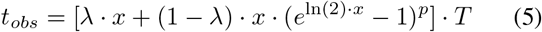

with *N* uniformly distributed points in *x* ∈ [0, 1], the total time length *T*, the linearity *λ* of the time spacing and the exponent *p* of the exponential time spacing.

A multi linear regression

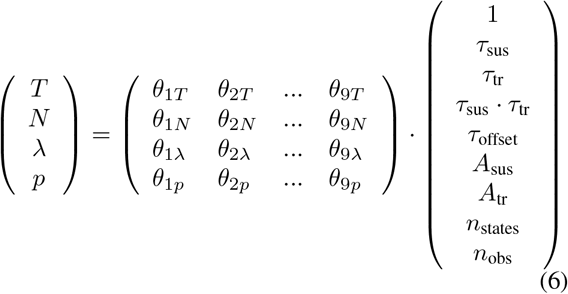

is performed between the time characteristics *N, T, λ* and *p* chosen by an experimenter, and the parameters given by the transient function *f*_tr_ that characterize the dynamics of the underlying system. The characterization of the dynamics allows to transfer typical experimentally chosen measurement times to other models. The time characterisitcs and the time predictors are on logarithmic scale except for the linearity *lambda* which lies between [0,1]. The multi linear regression is applied with intercepts *θ*_1*T*_, *θ*_1*N*_, *θ*_1*λ*_, and *θ*_1*p*_. The estimated regression coefficients 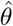 of Eq. 6 are shown in Supplementary Table 1. With 4 characteristics of the measurement times and 9 predictor variables, there are 36 regression parameters in total. The predicted time parameters of the total time length *T*, number of data points *N*, linearity *λ* of time spacing and exponent *p* of exponential time spacing for the 19 published models are shown in Supplementary Figure 1. The Pearson correlation coefficient *ρ* falls between 0.67 and 0.93. Note that our analysis has been performed to be independent on physical units by considering that the time scale of the model dynamics defines the time constants *τ* and thus, the time characteristics, independently of their units.

#### 2.3.3 Data simulation and measurement error

Because experimental data in molecular biology is typically log-normally distributed [23], the simulated data points

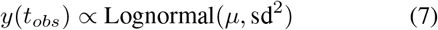

at the suggested observation times *t_obs_* are drawn from a log-normal distribution, where the mean *μ* is given by the expectation value of the model dynamics. Due to different experimental techniques and measurement methods, the inter-model variation of the measurement error is usually larger than the intra-model variation. Therefore, we assume a linear mixed-effects model on logarithmic scale to estimate the error parameters:

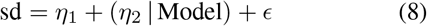

with *i* = 1, 2,…, *n*_obs_, a fixed effect *η*_1_, a model-dependent random effect 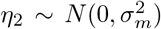 and a Gaussian noise 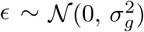. The estimated parameters for the 19 published models are *η*_1_ = −1.12, *σ_m_* = 0.25 and *σ_g_* = 0.015. An example of a realistic simulation of measurements is shown in Figure 2C and in Supplementary Figures 2 and 3.

**Figure 2:**
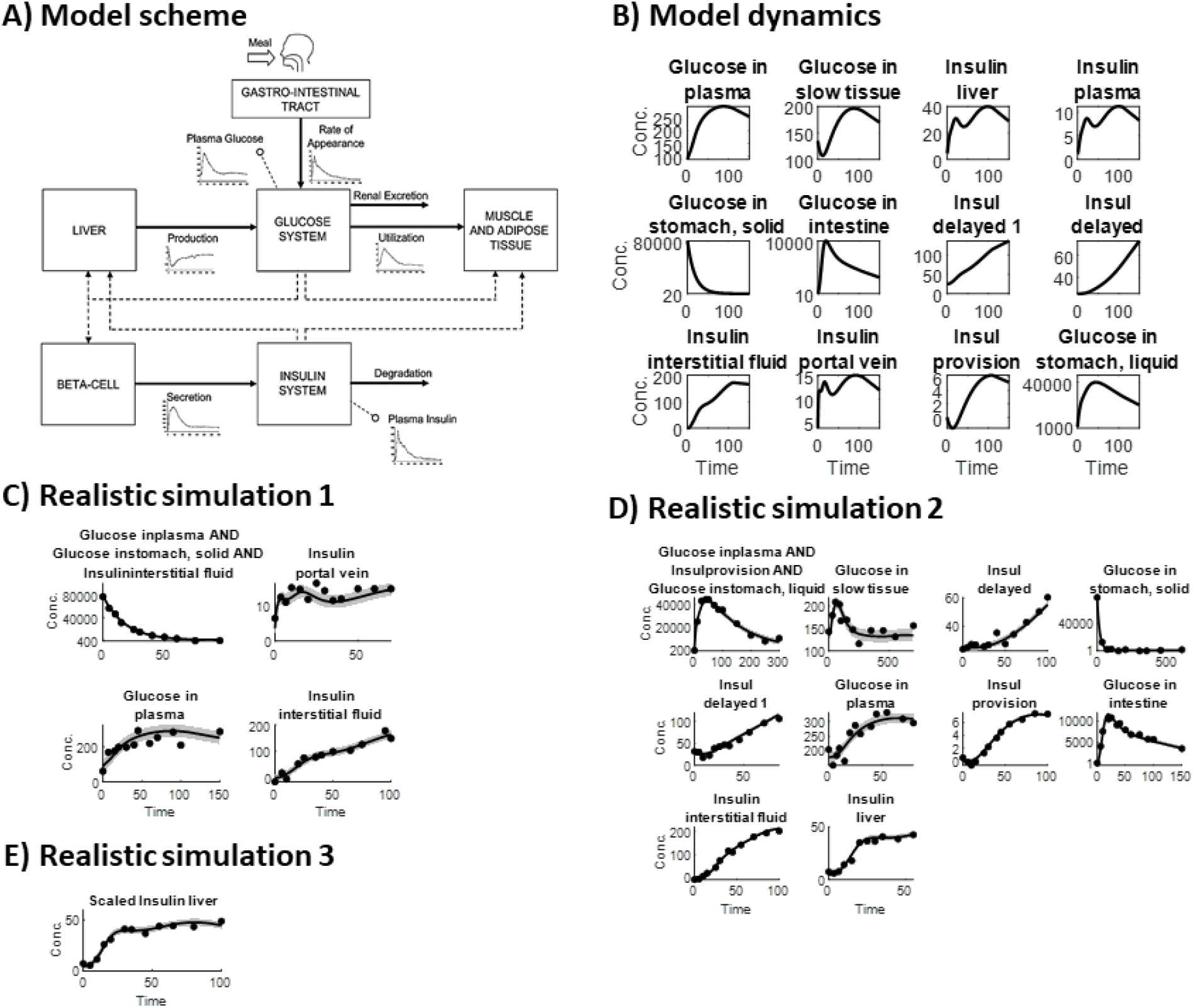
Example of creating a realistic design for a desired model. A) Model scheme of the glucose-insulin application example[8]. B) Model dynamics of the 12 glucose and insulin states. C) Simulation of a realistic measurement of the glucose-insulin model [8]. The model dynamics and simulated time and data points for the drawn observables are depicted.

### 2.4 Validation criteria

To validate how realistic the described simulation of measurements actually is, the realistic simulation is compared with a naive simulation, an unrealistic simulation and with the published model itself with the measured experimental data. The measurement characteristics of the naive and unrealistic simulation are displayed in Table 2.

**Table 2:** Measurement characteristics for the realistic, naive and unrealistic simulation. To validate how realistic the described implementation is, it is compared to a naive and unrealistic simulation with fixed numbers of directly measured observables, data points, and measurement error, with linear time spacing and naively chosen time duration.

| Measurement characteristics | Realistic design | Naive design | Unrealistic design |
| --- | --- | --- | --- |
| # Observables | Binomial draw [12-100] % | 50 % | 100 % |
| Observation type | $g^1, g^2, g^3$ | $g^1$ | $g^1$ |
| # Data points | Multi linear regression | 10 per obs. | 100 per obs. |
| Measurement error | Mixed effects model | 5 % | No error |
| Time spacing | Linear & exponential | Linear | Linear |
| Time duration | Multi linear regression | rounded to $10^n, n \in \mathcal{N}$ | Longest T of all obs. |

The simulations are compared to the published model using the root mean square deviation

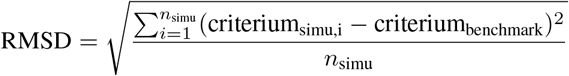

of 5 different validation criteria. The validation criteria are chosen to evaluate the parameter identifiability, integration of the ODEs, and parameter optimization performance of the model. For the least squares optimization

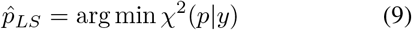

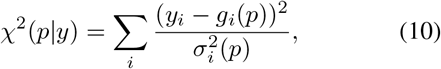

the number of fits *n*_optfits_ in the global optimum is evaluated by:

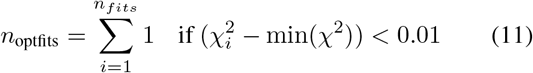

with *n_fits_* being the number of fits performed for multi-start optimization using different random initial guesses. To evaluate the identifiability, the number of non-identifiable parameters is assessed with the identifiability test by radial penalization [20]. Further, the number of function evaluations of the converged fits, the computation time for fitting and the number of steps for the ODE integration are compared.

### 2.5 Implementation

The algorithm for simulating realistic designs and experimental data is implemented in the MATLAB based modeling framework Data2Dynamics [28]. To import the model and export the realistic simulation in the standardized SBML [16] and PEtab [30] formats, the MATLAB code including the Data2Dynamics toolbox is as follows:

~~~
SBML2Model
arRealisticDesign
arExportPEtab
~~~

By default, the dynamic parameters of the ODE system are set to a logarithmic scale to improve the convergence of the optimization [29, 19] and the parameter bounds are set to four orders of magnitude.

## 3 Results

### 3.1 Application example

In the following, we demonstrate the general aspects of the algorithm on the ‘Meal Simulation Model of the Glucose-Insulin System’ by Dalla Man, Rizza and Cobelli, 2007 [8] which describes the physiology after meal uptake. It describes the concentration of insulin and glucose in different compartments. Here, glucose and insulin fluxes are taken from a human database. The model scheme and the 12 state dynamics of the glucose-insulin system are shown in Figure 2A and B.

According to the main steps of the implemented algorithm displayed in Figure 1, first, the number of observables and their observation type are set by a binomial draw according to Table 1 and as a subset of the given states. In the examples shown in Figure 2C and D, there are 4 and 10 observables each and one compound consisting of three states. Our approach for generating realistic designs can also draw compound observation function consisting of more states as illustrated in **??**E. In the third example of Figure 2E, only a single observable has been drawn which is measured on relative scale and, thus, has a scaling factor. After drawing the observables, the model is characterized for the observed dynamic states using the transient function (Eq. 4). With Eq. 4 an Eq. 6, the total time range *T*, the number of data points *N*, the linearity *λ* of the time spacing and the exponent *p* of the exponential time spacing are examined. The simulated measurement time points consist of *N* logarithmically spaced time values between *t*_0_ = 0 and *T*. The data is simulated at the suggested time points with a log-normal distribution whose mean is given by the model dynamics and the standard deviation is given by the error model (Eq. 8). Three examples of simulated data are shown in Figure 2C, D and E. Further application examples of a mitogen-activated protein-kinase (MAPK) cascade by Huang and Ferrell, 1996 [15] and of an Epidermal Growth Factor Receptor (EGFR) signaling by Kholodenko et al, 1999 [18] are shown in the Supplementary Figures 2 and 3.

### 3.2 Validation

Simulation studies for benchmarking should provide results that are as similar as possible to analyses based on real data. To validate how realistic the presented approach for the simulation of systems biology measurements is, we evaluate the realistic simulations on 9 published models. To ensure implementation and data consistency, the 9 models are selected such that they support the PEtab import in the Data2Dynamics toolbox. To assess the optimization and integration performance, we compared 50 realistic, 50 naive and 50 unrealistic designs against the published model by evaluating the 5 different validation criteria introduced in section 2.4 using 1000 multi start optimization runs for each model and simulation.

The performance of the 5 validation criteria for the realistic simulation, naive simulation, unrealistic simulation and the published model is shown in Figure 3. In most cases, the realistic simulation performs better than the naive and unrealistic designs, i.e. it is closer to the benchmark performance. For 8 of the 9 models, the median number of fits in the optimum of the realistic simulation is closest to the published model (all except Bruno [6]), and for 6 models also the RMSD is closest to the published model (green). In case of integration steps, the RMSD of the realistic simulation is the smallest only in 4 out of 9 models. However, in the remaining 5 cases, there are no major deviations, and the performance of the realistic simulation is comparable to that of the naive and unrealistic design. For the optimization process, the realistic simulation generally requires more function evaluations and more computation time than the naive and unrealistic design (downward trend, except for Fiedler [10] and Fujita [11]). Moreover, the number of function evaluations and the computation time depend strongly on the number of parameters. For instance, in case of the model by Crauste et al. [7], the number of parameters is highest for the unrealistic design, resulting in fewer converged fits and, thus, fewer function evaluations and a lower computation time. For the number of non-identifiable parameters (Figure 3) the realistic simulation deviates the most from the published model.

**Figure 3:**
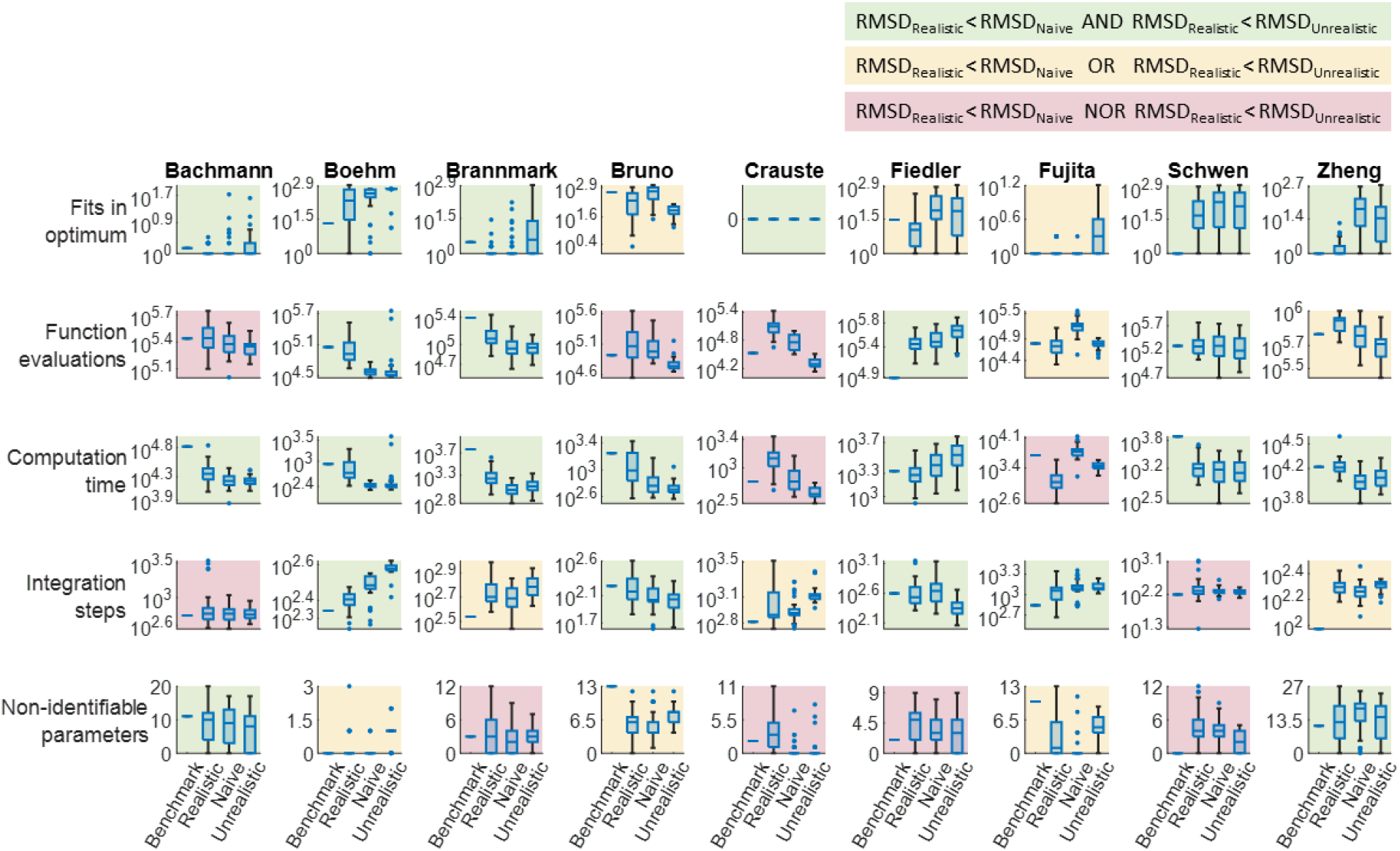
The optimization and integration performance for 9 published models, the realistic simulations of the 9 published models, and the comparison of naive and unrealistic simulations. Each boxplot comprises 50 simulations. To determine how realistic the simulations are, the optimization and integration performance should be close to the published model. This is illustrated by comparing the RMSD of the realistic simulation with the benchmark performance. If the realistic designs are closer to the published model in terms of the RMSD than the naive and unrealistic designs, the figure is highlighted in green. If the realistic designs are closer to the published model than the naive or the unrealistic designs, the figure is highlighted in yellow. If the realistic designs deviate more from the published model than the naive and unrealistic designs, the figure is highlighted red, which is the case for 11 of 45 comparisons.

## 4 Discussion

In this work we present an approach for creating realistic experimental designs and for simulating realistic data for any biological model. Many dynamic models of biological systems are publicly available, but lack the experimental data, which is crucial for evaluating and benchmarking of methods. Our approach fills this gap by providing a realistic setup for a given dynamic model, including observed quantities, observation time, number of data points, and measurement errors. This realistic simulation setup can be used as a basis for simulation studies to assess and improve methodological and computational procedures.

The analyzed models show large deviations in the number of observables, their observation type and also in estimating the observation time points with the standard error having the same order of magnitude as the mean. These variations are the key for the realistic diversity of biological experiments. By using a binomial draw for these measurement characteristics, the realistic simulations reflect this diversity in real applications. Still, the observation types of the observables depend strongly on the studied system and the possible measurement technique. The time duration, time spacing and number of data points for the suggested observables are set by the experimenter and thus vary between projects. Yet, the multivariate regression provides a good representation of these quantities.

The validation of the realistic simulation is challenging because the observables and measurement characteristics are drawn randomly and therefore vary widely, making the simulated experiments not directly comparable to the single realization of the published model.

The variety of designs in the published models described above actually calls for more than 19 examples to define typical experimental design. Also, more models would allow a better evaluation. In this work, we selected just one experimental condition for each model, i.e. the condition with the most species. Further studies with a more detailed representation could potentially improve our results. Possible refinements of a realistic model design are e.g. including repetition measurements, including multiple conditions such as genetic perturbations or stimulating with different doses, or a more representive selection of compound observables (which are more likely to be measured as a compound). Here, only a limited number of published models with data and comprehensive documentation how the data is linked to the model, have been available for the analysis of a realistic design. The realistic reflection of an experimental design obtained in this work are particularly promising for signaling pathway models. However, the results can also be applied to pharmacokinetic, curated, non-curated, metabolic, disease-related, or other dynamic models.

The results and algorithm presented in this study represent a realistic reflection of dynamic biological models. This realistic representation is essential for simulation studies to depict real-world applications and to evaluate methodological and computational issues in a realistic biological setting. The realistic simulation approach is also beneficial for benchmark studies because it takes away the freedom and dependence on the choice of simulation setup. resulting in less bias.

## Acknowledgments

This work was supported by the Federal Ministry of Education and Research of Germany [EA:Sys,FKZ031L0080 to J.E. and C.K.]; and the Excellence Initiative of the German Federal and State Governments [CIBSS-EXC-2189-2100249960-390939984 to C.K.].

