## Supplementary Material for "Realistic simulation of time-course measurements in systems biology"

### Supplementary material: Realistic simulation of time-course measurements in systems biology

JANINE EGERT CLEMENS KREUTZ*

#### Multi linear regression for the measurement characteristics

For a realistic choice of observation time points, multilinear regression with 9 predictor variables is performed to predict time characteristics such as the time duration $T$, the number of data points $N$, the linearity $\lambda$ of the time spacing, and the exponent $p$ of the exponential time spacing (Eq. 6). Model reduction is performed for each of the 4 time characteristics using the likelihood ratio test. Regression parameters that are not significant are marked with (-) in the Supplementary Table 1. All 4 time characteristics predicted by the multilinear regression show a strong positive correlation to the experimental measurement time characteristics observed in real data, with a Pearson correlation coefficient $\rho$ between 0.67 and 0.93 (Supplementary Figure 1).

#### Application examples

Supplementary Figure 2 shows additional examples for realistic designs of the Epidermal Growth Factor Receptor (EGFR) short term signaling by Kholodenko et al, 1999 [2] and Supplementary Figure 3 shows examples for realistic designs of the mitogen-activated protein kinase (MAPK) cascade by Huang and Ferrell, 1996 [1].

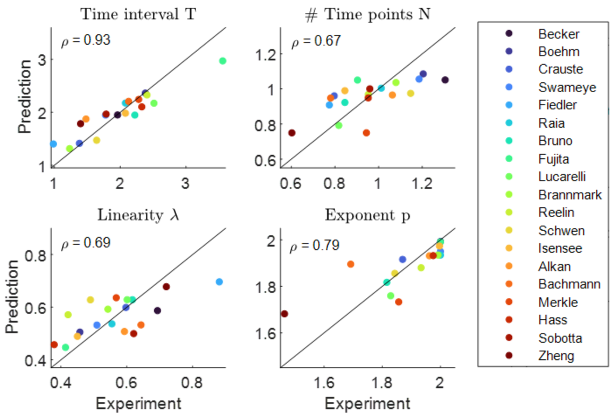


Supplementary Figure 1: The experimentally measured and the predicted time characteristics, of the overall time range $T$, the number of data points $N$, the linearity $\lambda$ of the time spacing and the exponent $p$ of the exponential time spacing for the 19 published models. A perfect match is indicated by the diagonal black line. The Pearson correlation coefficient $\rho$ is greater than 0.6 for all four time characteristics.

Supplementary Table 1: Estimated parameters $\hat{\theta}$ of the multilinear regression in Eq. 6 of the manuscript, for the overall time range $T$, the number of data points $N$, the linearity $\lambda$ of the time spacing and the exponent $p$ of the exponential time spacing and 9 predictor variables. Model reduction was performed using the likelihood ratio test, the parameters marked with (-) are not signifant.

| Predictor | $\hat{\theta}_1$ offset | $\hat{\theta}_2$ $\tau_{\text{sus}}$ | $\hat{\theta}_3$ $\tau_{\text{tr}}$ | $\hat{\theta}_4$ $\tau_{\text{sus}} \cdot \tau_{\text{tr}}$ | $\hat{\theta}_5$ $\tau_{\text{offset}}$ | $\hat{\theta}_6$ $A_{\text{sus}}$ | $\hat{\theta}_7$ $A_{\text{tr}}$ | $\hat{\theta}_8$ $n_{\text{states}}$ | $\hat{\theta}_9$ $n_{\text{obs}}$ |
|---|---|---|---|---|---|---|---|---|---|
| T | 1.58 | 0.13 | 0.28 | 0.042 | 0.00151 | - | -0.019 | -0.0046 | 0.016 |
| N | 1.02 | 0.037 | 0.059 | -0.03 | -0.00029 | - | 0.015 | 0.0016 | -0.014 |
| $\lambda$ | 0.616 | -0.014 | -0.044 | - | -0.00015 | -0.006 | -0.006 | -0.0039 | 0.0063 |
| $p$ | 1.73 | 0.08 | 0.098 | -0.035 | - | - | 0.031 | 0.0073 | -0.013 |

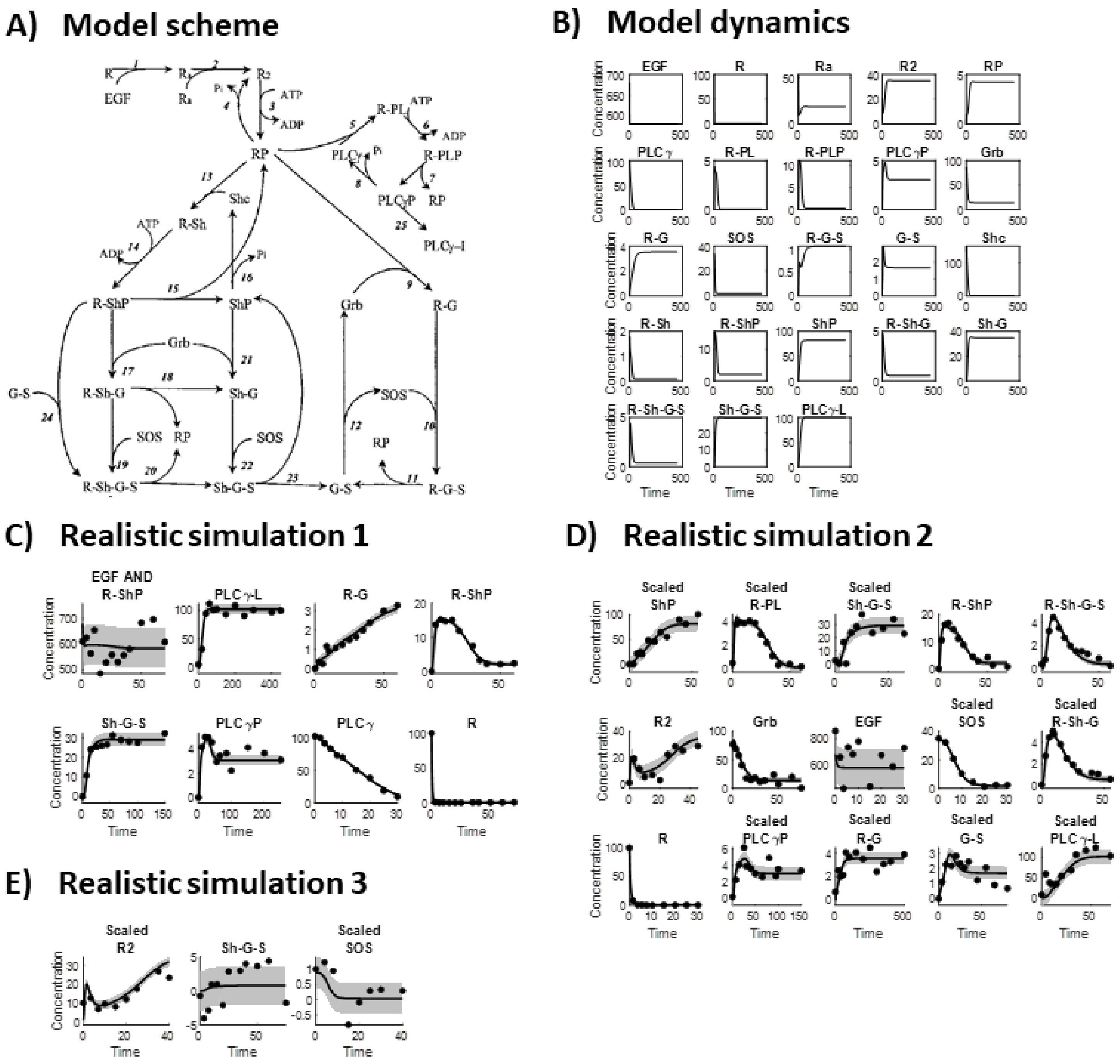


Supplementary Figure 2: Application example of an EGFR pathway model [2] using the realistic simulation approach presented in the manuscript. A) The model scheme of the EGFR short term signaling [1]. B) Model dynamics with the equations and parameters from the BioModels database. C) and D) and E) show three examples of data generated with different realistic simulation setups.

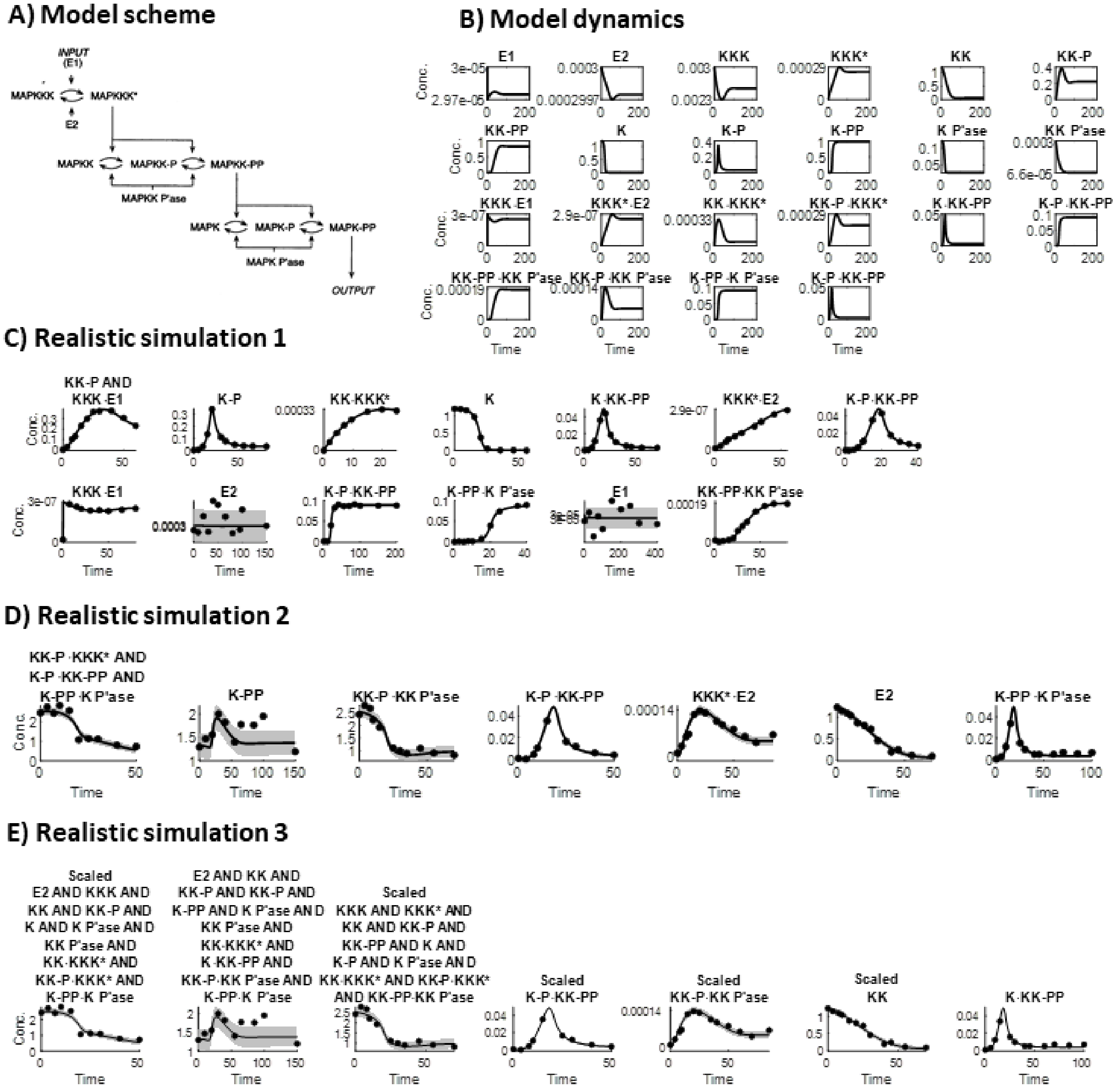


Supplementary Figure 3: Application example of a mitogen-activated protein-kinase (MAPK) cascade by Huang and Ferrell, 1996 [1] using the realistic simulation approach presented in the manuscript. A) The model scheme of the MAPK cascade [1]. B) Model dynamics with the equations and parameters from the BioModels database. C) and D) and E) Realistic simulations of measurements.
